# Evolutionary coexistence in a fluctuating environment by specialization on resource level

**DOI:** 10.1101/2021.05.18.444718

**Authors:** Meike T. Wortel

## Abstract

Microbial communities in fluctuating environments, such as the human gut or repeated dilutions in the laboratory, contain a wealth of diversity. Diversity contributes to the stability and function of communities and is maintained by underlying mechanisms. When nutrient levels fluctuate over time, one possibly relevant mechanism is that types specialize on low and high nutrient levels. The relevance of this process is supported by observations of coexistence in the laboratory, and by simple models, that show that negative frequency dependence of two such types can stabilize coexistence. However, as microbial populations are often large and numerous, they evolve. Our aim is to determine what happens when species can evolve; whether evolutionary branching can create diversity or whether evolution will destabilize coexistence.

We derive the selection coefficient in fluctuating environments and use adaptive dynamics to find that evolutionary stable coexistence requires a special type of trade-off between growth at low and high nutrients. We do not find support for the necessary trade-off in data available for the bacterium *Escherichia coli* and the yeast *Saccharomyces cerevisiae* on glucose. However, this type data is scarce, and might exist for other species or in different conditions. Moreover, we do find evidence for the right trade-off and evolutionarily stable coexistence of the two species together. Since we find this coexistence in the scarce data that is available, we predict that specialization on resource level is a relevant mechanism for species diversity in microbial communities in fluctuating environments in natural settings.

## 1 Introduction

Classical theory predicts that only as many species can coexist as there are resources^1,2^. However, in some environments, many more species coexist than there are resources^3,4,5^. A large body of theory explains this diversity from the use of different niches^6^ and the stabilization of diversity from frequency dependent interactions^7^.

In this study we focus on the diversity of micro-organisms in seemingly simple environments, with fluctuating resources. Species and strains can coexist over long timescales in complicated environments such as the human gut microbiota^8,9^. More surprisingly, even in laboratory environments, where the environment is simplified as much as possible, long-term coexistence of different strains is very widespread^10^. Several explanations have been given for this coexistence, including cross-feeding^11,12,13,14^, specialization on different resources^15,16^ and by specialization on resource level^17^.

The first observation of coexistence of two strains in an environment with a fluctuating resource level is by Levin^18^. With a mathematical toy model they showed that, when one species takes up nutrients faster at low resource levels and the other at high, coexistence is possible^17^. An extension of the model with fixed cycle lengths and lag phases also shows possibilities of coexistence^19^. In a different experiment, Turner et al. ^20^ observed coexistence and found evidence of a trade-off on growth at different resource levels, but a mathematical model based on Monod growth, described in the same article, showed that this was not enough to sustain coexistence. Closer analysis showed that indeed, in addition to the substrate specialization trade-off, there was cross-feeding between the strains. These studies are all restricted to analysing strains with fixed properties. To observe coexistence of cohabiting cells, the coexitence should have some evolutionary stability^21^, e.g. adaptation should not lead to the loss of the coexistence. Therefore we aim to study the relevance of specialization on resource level for evolutionary stable coexistence in fluctuating environments.

In this paper we present the selection coefficient in serial dilution experiments, which is the common way to do laboratory dilution and can therefore be directly applied. We supply a method to use experimental data together with computational model to obtain a explicit trade-off function. We show that certain trade-off functions can and some cannot lead to evolutionary stable coexistence. We combine the obtained trade-off data with the trade-off necessary for evolutionary coexistence and postulate that evolutionary coexistence is most likely in communities of two different species.

## 2 Model description

The model describes an environment where the resource is consumed, and at the moment the resource is completely consumed, cells are diluted to a fixed population size and new resource is added (for simulations a cut-off at very low substrate levels is used, after which the species densities hardly change). This relates to the conditions that are commonly used for experimental evolution. We focus only on the exponential and nutrient-limited growth phases and ignore lag-phase and stationary phase. We assume Monod-growth, because this captures the main features of growth at high and low substrate densities, and is also the most complex description for which we can derive an analytical equation of the selection pressure. Moreover, growth properties from more detailed models can be fit well to a Monod equations, making sure that our results will be confirmable with those models. This leads to the following dynamical system:

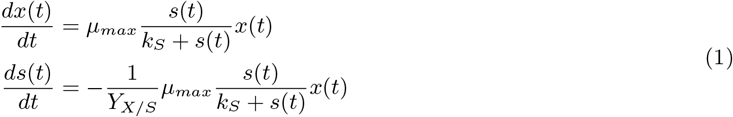

Where *x* is the biomass concentration, *s* the substrate concentration, *k_S_* the Monod constant and *μ_max_* the maximal growth rate. *Y_X/S_* is he numerical yield of the species x on the substrate *s*. To enhance readability for the derivations, we will use a notation where we write *s* for *s*(*t*), *x* for *x*(*t*), *μ* for *μ_max_, k* for *k_S_* and *y* for *Y_X/S_*.

## 3 Results

### 3.1 Selection in fluctuating environments

We calculate the selection coefficient *ω* as the ratio of the Malthusian parameters^22,23^ of the mutant and the resident minus one, such that positive values correspond to a mutant that can invade, and negative values to one that cannot (see Appendix Section A.2). We obtain the following selection coefficient:

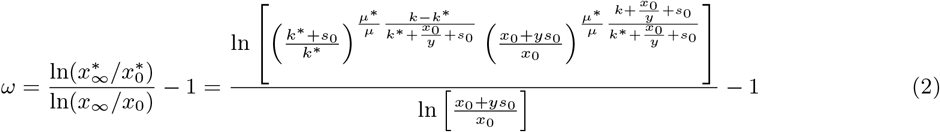

where mutant trait values are denoted with an *. This selection coefficient can directly be applied to experimental design and replace numerical simulations to estimate the relative selection on different growth traits^24^. Some direct observations from the selection coefficient are that there is no selection on the numerical yield, although the resident yield does influence whether other types can invade, and that higher initial substrate leads to stronger selection on the maximal growth rate, while higher initial biomass leads to stronger selection on the Monod constant. The derivation of this equation, calculation of the selection gradient and more insights on the parameter dependencies can be found in the Appendix Sections A.3 and A.5.

### 3.2 Evolutionary coexistence

Stable coexistence requires frequency dependent selection, i.e. fitness increases with decreasing frequency, which arises when one type specialises on high resource and reduces the high resource phase when abundant, making the environment relatively more beneficial for a type specialising on low resource and vice versa (Figure 1B-D). This cab kead to stable coexistence of the two types as long as there are no evolutionary changes, which we will further refer to as ecological coexistence ^21^.

**Figure 1:**
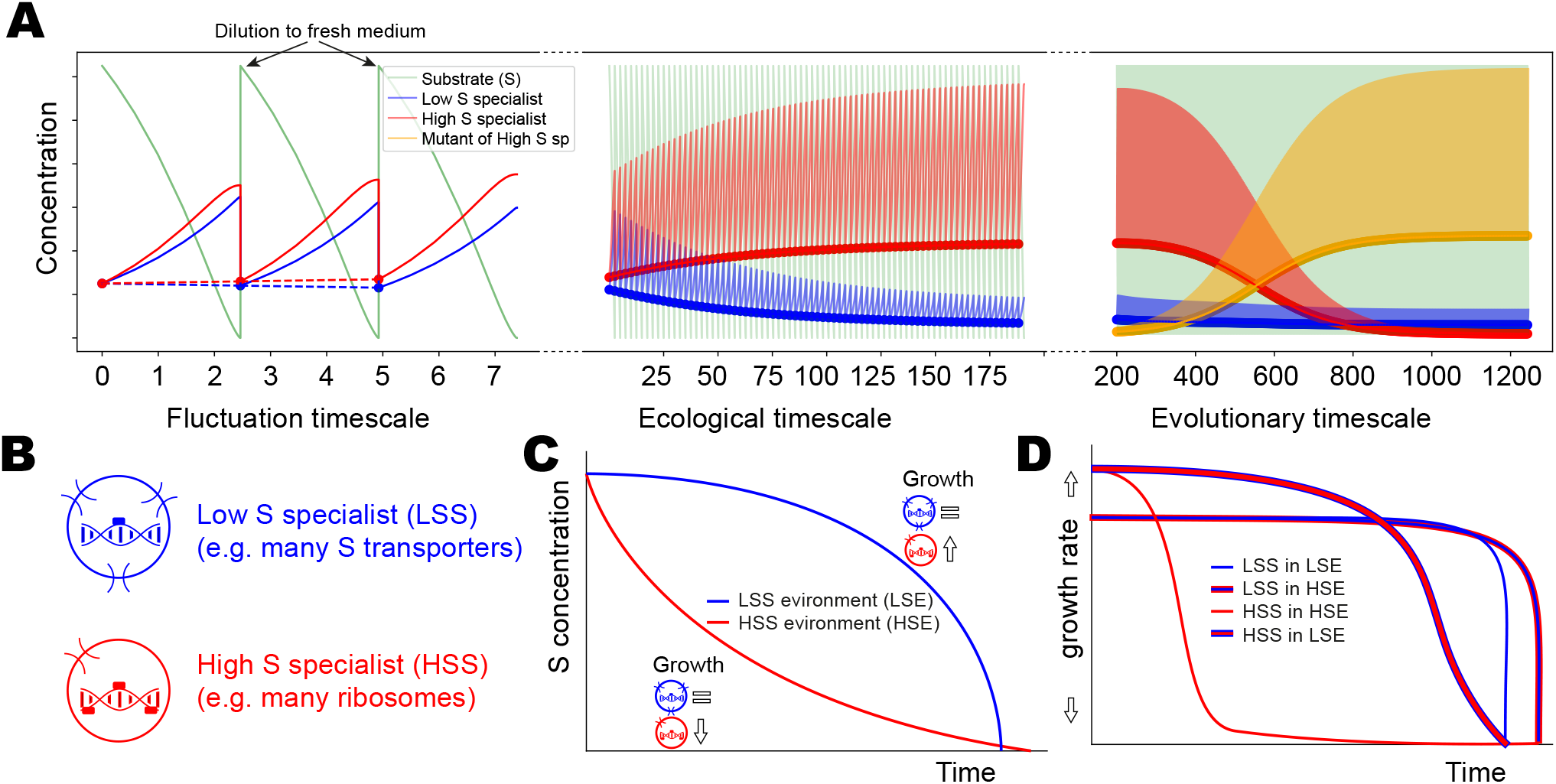
**A** The dynamics of serial transfer experiments. On fast ‘fluctuation timescales’, the species densities and metabolite concentrations are continuously changing. A stable pattern can arise over ecological timescales, where the concentrations at the beginning of each transfer remain constant. Over evolutionary timescales, strains can evolve properties such as maximal growth rate and affinity. Invading mutants either displace one of the strains and form a new ‘ecologically stable’ coexistence (as in the figure) or take over the whole community. **B** Negative frequency dependent selection between low and high substrate (S) specialists (arising from e.g. different protein allocations; investment in ribosomes for higher *μ_max_* and in transport protein for higher affinity (lower *k_S_*)).
**C** When low substrate specialists (LSS) are abundant, the substrate concentration decreases slower but is completely depleted faster than when high substrate specialists (HSS) are abundant (blue vs red line). LSS are not influenced greatly by the substrate concentration, but HSS thrive under the high substrate concentrations that are around for longer when LSS are abundant. Therefore HSS keep a higher growth rate for longer in an environment with LSS (red line with blue edges in **D**), than in an environment with HSS (red line in right panel). Similarly, LSS keep growing longer in an environment with HSS, because it takes longer for the substrate to be completely depleted (blue line with red edges vs blue line).

We study evolutionary changes, which could destabilize ecological coexistence, with evolutionary stable strategies (ESSes), i.e. a strategy (*μ, k, y*) that cannot be invaded by any other strategy 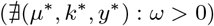. ESSes arise then the properties (*μ_max_* and *k_S_*) are bounded or correlated, e.g. high μ are associated with low *K_S_*(trade-offs as in Figure 2A and C). Trade-offs can lead to evolutionarily stable strategies (Figure 2B) or evolutionary singular coalitions (a state with more than one phenotype that cannot be invaded; e.g. Figure 2D)^26^. When the coexistence is stable even under evolutionary changes, we define it as evolutionary coexistence (termed ‘evolutionarily stable coexistence’ by Li et al. ^27^).

**Figure 2:**
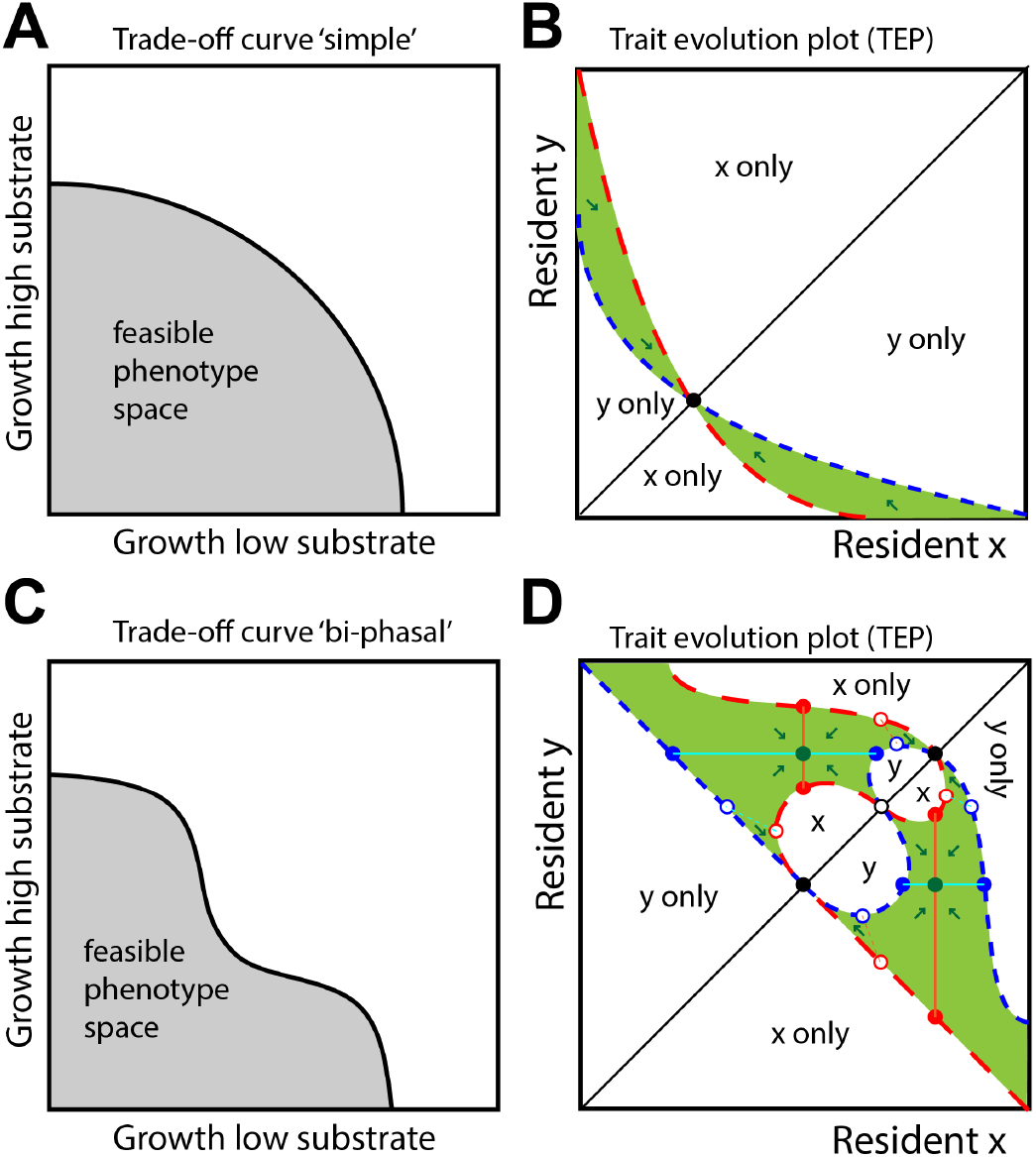
Schematic representation of different trade-offs and how they affect evolutionary coexistence. Arrows indicate selections gradients or evolutionary dynamics (graphical derivations of arrows in Appendix Figure A.10^25^). **A** When there is a more straightforward trade-off (‘simple’ trade-off), the evolutionary dynamics in the green coexistence area (**B**) go towards the point where both residents are the same, i.e. coexistence is lost over evolutionary time. **C** When there is a more complicated trade-off, for example this ‘bi-phasal’ trade-off, it is possible that there is an attracting point in the interior of the coexistence area (green dot in **D**), i.e. coexistence is evolutionarily stable.

In adaptive dynamics ecological and evolutionary timescales are separated. In this system, the ecological timescale coincides with the dynamics over the course of many transfers, the evolutionary timescales and the evolutionary timescales with the introduction of new types with new properties. Our system includes a third timescale, the fluctuation timescale, which encompasses the changes within a transfer (see Figure 1A). This extra timescale makes the ecological steady state a stable fluctuation, instead of a steady state, as is more common in adaptive dynamics.

Using the selection coefficient we can investigate the evolutionary dynamics using Trait Evolution Plots^26,28,25,29^. When the trade-off between maximal growth rate and affinity is ‘simple’ (Fig. 2A), there is only a small area of mutual invasibility between two strains with different properties (2B). In this area there is ecological coexistence, but, as the arrows show, evolution will destabilize this coexistence and the evolutionary outcome will be a single stable strategy (see Appendix Figure A.10 for the derivation of the arrows). However, a more ‘biphasal’ trade-off (Fig. 2C) can lead to a larger area of mutual invasibility with an evolutionary stable coalition (Fig. 2D and Appendix Figure A.10). There is no evolutionary branching, which means that this coexistence cannot arise by small mutational steps from a single ancestor, and has to arise from large mutational steps or by migration.

### 3.3 Case study: Trade-off in growth parameters for *Escherichia Coli* and *Saccharomyces Cerevisiae*

We have used information about *E. coli* and *S. cerevisiae* to study whether the trade-offs between growth at different resourde levels of organisms fall in the category of a ‘simple’ trade-off or a ‘bi-phasal’ trade-off, and to study the resulting evolutionary dynamics. We have collected experimental data for *E. coli* ^30^ and *S. cerevisiae* ^31^. Since the direct experimental data is limited, we have found a method to derive the trade-off from computational models. For *E. coli*, we used a mechanistic computational model of central carbon metabolism, studied the growth curves at a range of substrate concentrations for all paths through the metabolic network, fitted those curves with the Monod equation and then fitted a curve through the edge of phenotype space (the set of *μ_max_* and *k_S_*) to obtain the trade-off curve (see Appendix Figure A.1)^32^. For *S. cerevisiae* we used a more coarse-grained metabolic model, which we optimised at different glucose concentrations. Then we studied the growth curves of the different optimized models at a range of substrate concentrations, fitted them to the Monod equation and fitted a curve through the obtained set of *μ_max_* and *k_S_* to obtain the trade-off curve (see Appendix Figure A.2)^33^. In some cases the trade-off is related to transport kinetics^31^, but mostly it is a property of the whole cell and surfaces even with a single type of transporter. The trade-offs for the same species resulting from different sources leads to different trade-offs (see Figure 3). However, the shape of the curves do show some consistency, and resemble the ‘simple’ trade-off (Figure 2A). See Appendix A.4 for details on the fitting of the trade-off curves and the calculations of the numerical yields.

**Figure 3:**
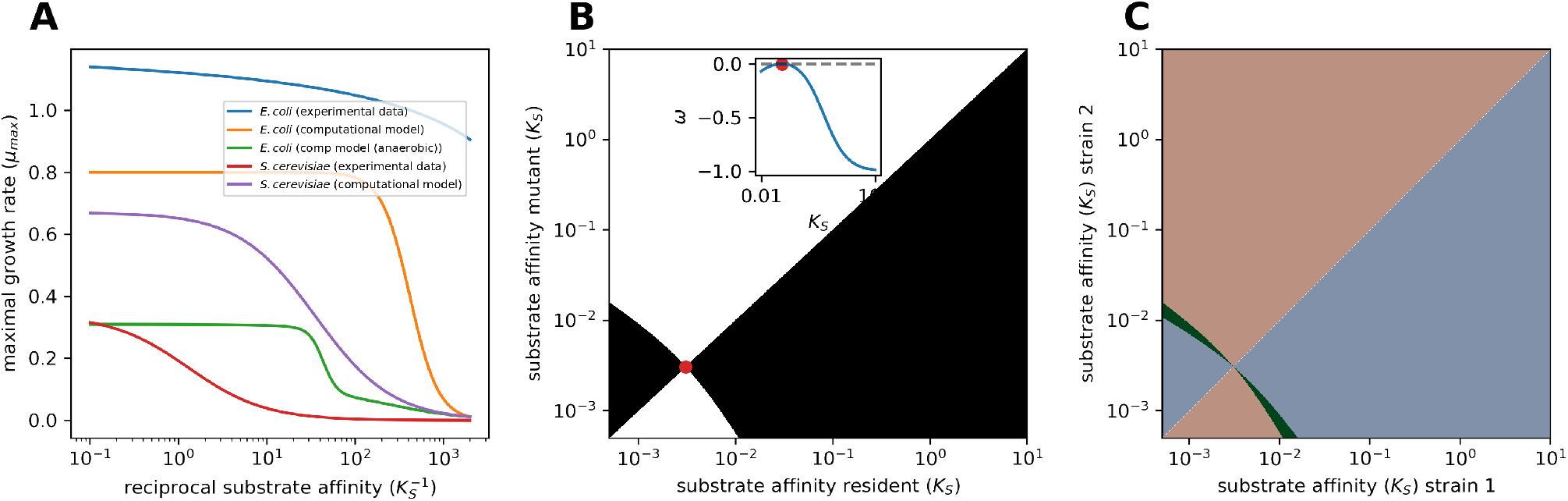
Trade-offs between growth parameters. **A** Trade-off between (*μ_max_*) and the reciprocal of the affinity 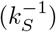. **B** Pairwise invasibility plot for the experimentally derived trade-off for *E. coli* at the conditions of the LTEE (initial glucose = 0.1375 mM, initial cells are 0.5·10^6^ cells/ml)^24^. The inset shows that the equilibrium is invasion stable. The equilibrium is also attracting and therefore an evolutionarily stable strategy (ESS). **C** Mutual invasibility plot. By reflecting the PIP of **B** over the main diagonal and overlaying it with the PIP, we can determine the area of mutual invasibility and therefore ecological coexistence (green area).^26^

Next, we use the trade-off to construct the pairwise invasibility plot for *E. coli* under conditions commonly used for repeated transfer experiment (Figure 3B). A single ESS emerges; if the resident would be very adapted to low substrate (low *K_S_*), higher substrate specialists can invade, and if the resident is adapted to higher substrate (high *K_S_*), lower substrate specialists can invade. In the ESS, no other type can invade (inset in Figure 3B). We can overlay this PIP with the PIP reflected in the main diagonal, to see when two types can mutually invade (green area in 3C), showing the area of ecological coexistence.

The obtained analytical expression for the selection coefficient allows us to investigate many different scenarios. However, for all trade-offs, from experimental and computational data, and for different initial substrate and biomass concentrations, the general picture is similar (see Appendix Section A.6): There is a small area of ecological co-existence, but analysis of the shape of the mutual invasibility plot shows that this coexistence is not evolutionary stable and the only attracting point is a single-type ESS. This means that two strains adapted to different conditions, such as a high-nutrient and low-nutrient condition, could coexist when placed together in a fluctuating environment. However, when they adapt in the same fluctuating regime, they will evolve in such a way that one of the strains will be lost, and a single strain will remain.

### 3.4 Evolutionary coexistence in multiple species consortia

One way to obtain ‘bi-phasal’ phenotype space, is to look at the evolutionary dynamics of two coexisting species. The methods we use are not adapted for multiple species. However, we found a simplification, which allows us to study evolutionary coexistence analytically, thereby being able to check many different scenarios, while still guaranteeing that the found solutions will also be a solution of the complete two-species dynamical system. We simplified the two trade-offs, because a species with a higher affinity or higher maximal growth rate for the same value of the other property, will always out-compete the other. As a proof of principle, we combine the trade-offs of *E. coli* and *S. cerevisiae* under anaerobic conditions, and will use the computational trade-off for *E. coli* and the experimental trade-off for *S. cerevisiae*. Although this is an artificial selection (different type of data and different conditions) of trade-off curves (solely based on their intersection), this will illustrate the method and its applicability for biologically realistic estimates as contrasted to a complete toy model. First we construct a new trade-off that combines the two (see Figure 4A and Appendix Section A.4.7). As species cannot convert into another one by evolutionary change at the studied timescale, we discard mutations that cross the intersection of the trade-off curves. However, since this intersection is in a fitness minimum, such crossings is almost guaranteed not to happen, and no additional constraints have to be imposed.

**Figure 4:**
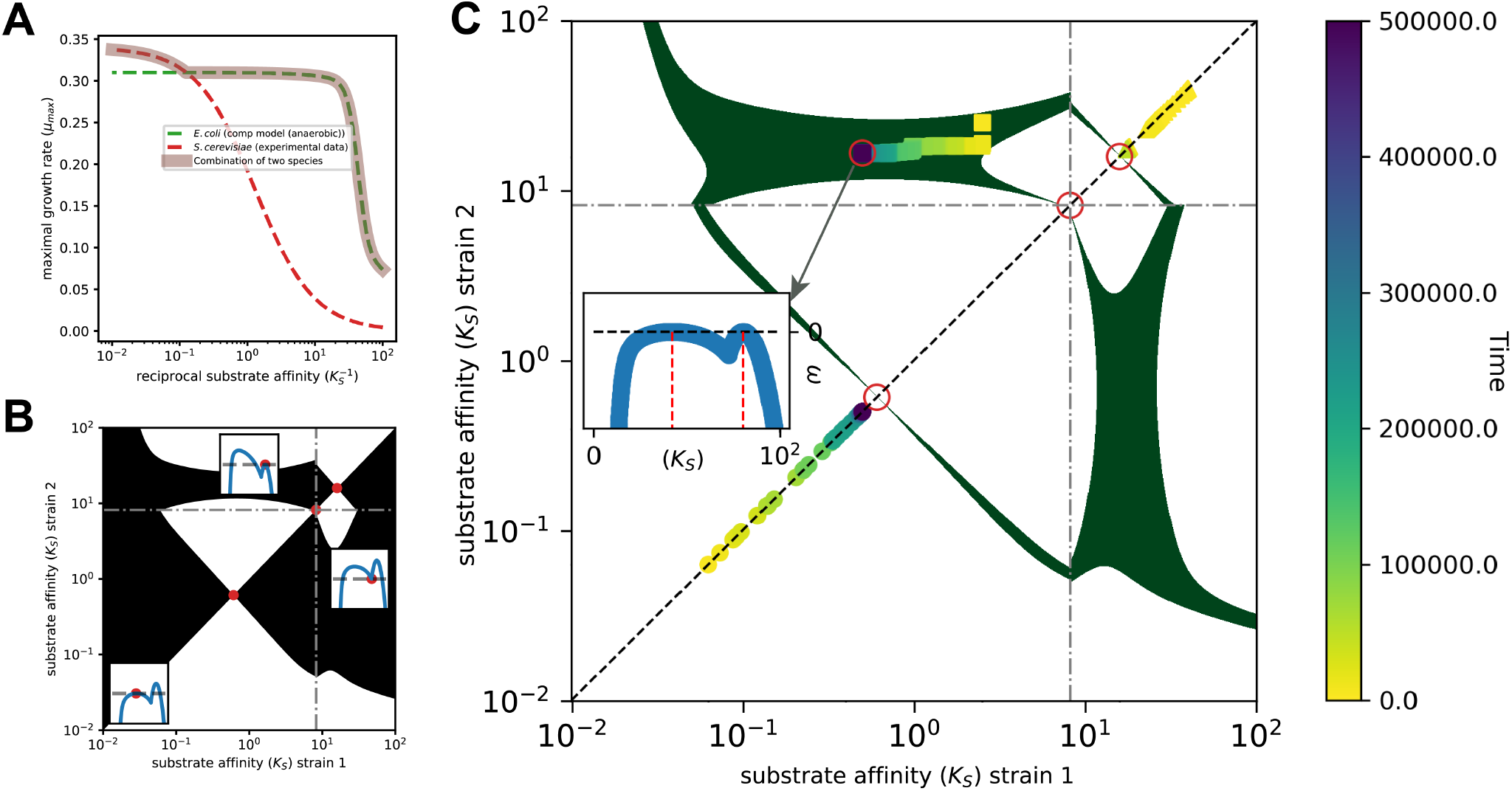
Combined trade-off of two species. **A** Trade-off between (*μ_max_*) and the reciprocal of the affinity 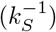. **B** In the PIP (with *x*_0_ = 10^6^ cells ml^-1^ and *s*_0_ = 500 mM) we can see there are three equilibria, of which the middle one is a local minimum corresponding to the switching point of one species to the other (gray dashes lines; those cannot be crossed by mutations). There are two equilibria that are attracting and locally invasion stable, which correspond to the ESS for each species. **C** Three different simulations are plotted onto the area of coexistence, created with the mutual invasibility plot (see Appendix Figure A.11 for the simulations). When initially one species is present, a single species ESS is reached. However, when we start with two species within the area of coexistence, an evolutionarily singular coalition (ESC) is reached. As shown in the inset, when in this ESC all other types have a negative selection coefficient, and therefore this ESC is evolutionarily stable.

The analytical approach allows us to screen different conditions (initial substrate and biomass concentrations) and find scenarios of coexistence (trait evolution plots resembling Figure 2D). When we construct the pairwise invasibility plot for the two-species trade-off, there are three equilibria, one local minimum at the species intersection and two local maxima, one for each species (Figure 4B). The single species maxima are both ESSes and will be reached when we start with a single species (Figure 4C and Appendix Figure A.11). There is an area of mutual invasibility between the species, shown as the green area in Figure 4C. If both species would be adapted alone under these conditions, they would coexist when one migrates into the other community. Moreover, using the analysis of trait evolution plots (Appendix Figure A.10), we can show that there is an evolutionary attracting point in the species coexistence area^25^. With simulations (Appendix Figure A.11) we can find this point and we show numerically that in this point, the selection coefficient for all other types is negative (inset in Figure 4C). This means that after the species coexist together, they will continue to evolve, but reach an evolutionary singular coalition.

## 4 Discussion

Nutrient fluctuations are abundant in nature, and are often mimicked in laboratory evolution studies with a serial transfer regime. In this paper we derive the selection coefficient in such regimes, develop a method to use computational models to obtain trade-off curves between growth parameters and apply adaptive dynamics to study the evolutionary dynamics. From the limited available data we do not find any evidence for evolutionary stable coexistence by specialization on resource level, making this less likely to be an explanation for the observed diversy in laboratory evolution studies^10^. However, it could be a factor contributing to co-existence, such as in Turner et al. ^20^, where this specialization is combined with cross-feeding. We show with a proof of principle that the methods can in this case be applied to species coexistence, and although we do not have data on naturally co-occurring species, from our results we postulate that evolutionarily stable coexistence by specialization on resource level is much more likely to occur.

Although we have not been able to show it rigourously, the evolutionary stable coexistence depends on the shape of the trade-off curve (see Figure 2). Combining the two species leads to a more ‘bimodal’ trade-off curve (with two ‘bumps’, see Fig. 4). In theory, a single species could have such a shape trade-off curve. We have not found any evidence for this, but as our data here is limited, we do not exclude the possibility. An interesting future direction would be investigate if it would be possible to determine the existence of evolutionarily stable coexistence from the shape of the trade-off curves. Perhaps the methods used in Ehrlich et al. ^34^ could be a start for finding such results. The distinction between species and strains is made here on the bases of the tradeoff curves. Strains are variants that are evolutionarily constrained by the same trade-off curve, while different species follow different trade-off curves. This could be interpreted as species being further apart in phenotype space than strains. We do acknowledge that other properties will also affect the trade-off curve, and trade-off curves themselves can evolve over longer timescales^35^.

Our results give indication which conditions are most promising to lead to coexistence. Evolutionary coexistence between species is most likely when dilution is not too extreme, and relatively high cell densities remain before nutrients are replenished. This is because high dilution leads to a long growth phase, which benefits a fast growth strategy, making it difficult for low substrate specialists to survive (this was also found in Abreu et al. ^36^). Natural habitats such as the mammalian gut satisfy such conditions.

We have used a trade-off between maximal growth rate and substrate affinity to determine the possible trait space of the organisms. Data and detailed models that can infer these properties are not abundant, therefore we focussed on *E. coli* and *S. cerevisiae*, where data is still limited and not completely consistent (as can be observed from the difference between the curves obtained from different datasets in Fig. 3A, indicating that more study is needed on these trade-offs.). We accept that these might not be the most biologically relevant species for coexistence studies, but if we can already find possibilities for coexistence in this very limited dataset, it is likely to occur for some combinations that cohabit in nature. We have shown some support for the trade-off in these species, however Velicer and Lenski^37^ set out to measure this trade-off experimentally in *E. coli* and found more contra-than supporting evidence. However, as they also note, perhaps their experiment was not long enough to actually reach the trade-off, and general growth performance mutations on the supplied substrate were selected for. Moreover, *E. coli* might not be the organisms where this phenomenon is most important, because it tends to use active transport that usually already coincides with a high affinity. This trade-off might be more relevant for other micro-organisms, for CO2 uptake by cyanobacteria and green algae^38^ or plants (see supplementary information in Beardmore et al. ^39^).

Our basic model uses Monod growth, which is a rather simple description of microbial growth. However, it is also the most complicated one for which we can derive an analytical expression for the invasion fitness, which is a new result obtained here. With more detailed models we can do similar analyses, although we have to resort to simulations to obtain the invasion fitness. However, for the more detailed models used to derive the trade-offs in this paper^32,33^, we do expect to find the same results, because the important properties are captured in the fit of the trade-off curve. The only differences would arise from the error of the fit and from the way we would include mutations, as we have only studied adaptation on the pareto front, i.e. with strategies optimal in the *μ_max_-k_S_* domain. This does lead to two interesting directions for further investigation: Simulation with more detailed growth models, from a Hill equation to a metabolic network or a genome scale model with uptake kinetics^40^, and including mutants that are not directly perfectly adapted, which could be done with (agent based) simulations (e.g. Virtual Microbes^41^).

In Figure 4C we see bi-stability in the evolutionary sense, i.e. the reached ESS depends on the initial traits of the species. However, the proposed model can, besides coexistence, also show ecological bi-stability, i.e. the final outcome depends on the initial ratio of the strains. Bistability was also observed in another study with a trade-off between maximal growth rate and lag phase^42^. The conditions for bi-stability of single-strain states include mutual non-invasibility, and can therefore be studied with Equation 2. Bi-stability is only possible if the strain specialized on high substrate has a low numerical yield, such that when that strain is abundant, nutrients remain high, favoring itself and supplying the necessary positive feedback. However, when we look at the measured parameters, bi-stability is not observed (there is no overlay of non-invasibility areas in figure A.9). Note that bi-stability is not possible when instead of the initial biomass, the dilution is kept constant, because then the yield does not affect the invasion fitness anymore (see Appendix Section A.3.1).

We have only discussed the co-existence of two strains or species. Previous studies have shown that theoretically, more than two species could coexist, albeit mostly though numerical simulations or only showing neutral coexistence^17,19,42^. Based on trade-offs implemented here, it is unlikely for more than two strains to stably coexist solely on resource level specialization. Still, if specialization on an intermediate level of resource is a side-effect, it can still help a new variant to (temporally) gain some benefit. In this paper we have studied the effect of resource level specialization in isolation, to conclude how strong the effect of only this aspect can be. However, the interplay between resource level specialization and other properties could play a role in maintaining community diversity, and as a future direction it would be interesting to study more complex communities (e.g. ^43,44^) and see if resource level specialisation plays a role.

Several open question remain. First, we have modeled predictable transfers after complete substrate depletion. It would be interesting to study whether less regular fluctuations influence the results. If nutrients are not completely depleted, this will benefit the high substrate specialist, and therefore coexistence will be less likely, but the conditions for coexistence could be investigated. Second, in the coexistence, the high substrate specialist is often more abundant than the low substrate specialist. If there would be a relation between the equilibrium ratio for the different strategies and the location of the ESC, this could simplify finding the ESC, which is difficult because an analytical solution for the fitness with more than one resident cannot be found. However, we have not been able to find this relation. Third, it would be interesting to apply our methods to other growth and community dynamics, such as lag phases, death in stationary phase and cross-feeding. It might be possible to assign the relative contributions to coexistence when combining different properties in one model. Last, it would be ideal to have experimental verification of these results. For that, first, models and theory can be used to identify an experimental system that would be suitable, i.e. where we expect evolutionarily stable coexistence under some (initial) conditions. Then, experiments can be designed to determine if only resource level specialization achieves coexistence, or if it is one of the contributors. As long as a suitable experimental system is lacking, virtual organisms could be a first test.

In conclusion, we have shown that we can use adaptive dynamics and trade-offs based on experimental data and computational models to show the biologically relevance of an observed and suggested mechanism behind diversity of microbial communities in fluctuating environments. As more quantitative data becomes available and more mechanistic models can be parameterized, we can investigate how common it is that species could coexist by specialization on resource level. Moreover, we supply the selection coefficient in fluctuating environments and techniques to investigate evolutionary stable diversity within and between species, which can be applied more broadly.

## 5 Acknowledgements

I would like to thank Han Peters for help with the mathematical derivations, Hans Metz for help with applying adaptive dynamics techniques, Guilhem Doulcier for the online availability of his lecture notes and code on adaptive dynamics and Juan Bonachela, Bob Planqué, Martijn Egas and Huub Hoefsloot for helpful discussions.

## Notes

### Competing Interest Statement

The authors have declared no competing interest.

### Summary of Updates

Included analysis of the evolutionary dynamics in the trait evolution plots, shortened (Fig. 2 and 3 combined, Fig. 4 and 5 combined), clarified (new Fig. 2) and updated the supplemental information.

