## Appendix for "Evolutionary coexistence in a fluctuating environment by specialization on resource level"

Meike T. Wortel[1,*]

[1]Molecular Microbiology and Microbial Food Safety, Swammerdam Institute for Life Sciences, University of Amsterdam, Amsterdam, The Netherlands
[*]

### A Appendix

#### A.1 Parameters and variables

| Symbol | Name | Units |
|---|---|---|
| $x$ | Biomass | $10^6$ cells ml$^{-1}$ |
| $s$ | Substrate | mM |
| $\mu$ or $\mu_{max}$ | Maximal growth rate | h$^{-1}$ |
| $k$ or $k_S$ | Monod constant (substrate at half maximal growth) | mM |
| $y$ or $Y_{X/S}$ | Yield of biomass on substrate | $10^9$ cells mmol$^{-1}$ |

#### A.2 Growth conditions, fitness and selection gradient

We study the conditions where periodically substrate is added to the system, and cells are diluted to a fixed concentration. To focus solely on the growth phases, we study the system until the substrate runs out, and we stop when $s(t)$ approaches 0. This is equivalent to transferring at long enough timescales such that all substrate is consumed, when there is no death at stationary phase. In some conditions, death at stationary phase is observed, and trade-offs between death rate and for example maximal growth rate are possible, but here we want to isolate the effects of only the growth phases. The dynamical system is:

$$\begin{aligned} \frac{dx}{dt} &= \mu \frac{s}{k+s} x \\ \frac{ds}{dt} &= -\frac{1}{y} \mu \frac{s}{k+s} x \end{aligned} \tag{A.1}$$

Where $x$ is the biomass, $s$ the substrate, $k$ the Monod constant, $\mu$ the maximal growth rate and $y$ the numerical yield.

Initially, we start every cycle of substrate addition with $s(0) = s_0$ and $x(0) = x_0$. Selection here acts on the increase of growth over a full cycle. We therefore define the final concentration divided by the initial concentration

$(x_\infty/x_0)$ as the growth factor. For the situation with only one type, at the end of the growth $s(t = end) = 0$ and $x(t = end) = x_0 + ys_0$ and therefore the growth factor is equal to $\frac{x_0+ys_0}{x_0}$.

When we define the invasion fitness, we have to assess the growth factor of a mutant ($x^*$) with a different Monod constant ($k^*$), maximal growth rate ($\mu^*$) and/or yield ($y^*$) in the dynamic conditions set by the resident strain (Equations (A.1)). If the mutant has a higher growth factor than the resident ($x$), the mutant increases in relative abundance and will be able to invade. More precisely, a mutant can invade when:

$$\frac{x^*(\infty)}{x^*(0)} > \frac{x(\infty)}{x(0)} \tag{A.2}$$

We define the fitness ($W$) of the mutant as the fraction of the Malthusian parameters[1,2]:

$$W = \frac{\log(x^*_\infty/x^*_0)}{\log(x_\infty/x_0)} \tag{A.3}$$

The log (throughout this indicates the natural logatrithm) is commonly used to compensate for exponential growth, but does not affect the fact that strains with $W > 1$ can invade. In evolutionary analysis, the selection coefficient is used. The selection coefficient $\omega$ (commenly called $s$ but here denoted with $\omega$ to avoid confusion with the substrate concentration) is defined as $W - 1$ and therefore a selection coefficient of 0 means that a mutant is not selected for or against (the same results can be obtained using the selection-rate constant $r = \log(x^*_\infty/x_\infty) - \log(x^*_0/x_0)$[1,3,4]). Strains with positive selection coefficients in a resident population will increase in number when rare, and can invade.

By studying the effect of the different phenotypes (in our case $\mu$, $k$ and $y$) on the fitness, we can calculate the selection gradient (see Appendix Section A.5). The definition of the selection gradient ($S_X$) of a property $X$ is (as in[5]):

$$S_X = \frac{X}{W}\frac{\partial W}{\partial X} \tag{A.4}$$

#### A.3 Derivation of the mutant fitness and the selection gradients

To find a fitness of a mutant ($x^*$), we need to calculate how it increases in frequency when it invades in the population of a resident ($x$). We will assume that this mutant is in such a small amount that the mutant does not affect the metabolite profile $s(t)$. Then, we will deduce the growth factor ($\frac{x^*(\infty)}{x^*(0)}$) for the mutant.

$$\begin{aligned} \frac{dx^*}{dt} &= \mu^* \frac{s}{k^*+s} x^* \\ \frac{1}{x^*}\frac{dx^*}{dt} &= \mu^* \frac{s}{k^*+s} \\ \frac{d}{dt}(\log x^*) &= \mu^* \frac{s}{k^*+s} \end{aligned} \tag{A.5}$$

Note that the right hand side depends on $t$ but not on $x^*$, because we assumed that the metabolite profile is determined by the resident. Thus:

$$\begin{aligned} \log(x^*(\infty)) - \log(x^*(0)) &= \int_0^\infty \mu^* \frac{s}{k^*+s} dt \\ \frac{x^*(\infty)}{x^*_0} &= e^{\int_0^\infty \mu^* \frac{s}{k^*+s} dt} \end{aligned} \tag{A.6}$$

To calculate the above integral we change variables from $s(t)$ to $s$. Because we know that $s$ runs from $s_0$ to 0, we integrate from $s$ from $s_0$ to 0 in stead of $t$ from 0 to $\infty$. We do have to multiply with the derivative $\frac{dt}{ds}$:

$$\int_0^\infty \mu^* \frac{s(t)}{k^*+s(t)} dt = \int_{s_0}^0 \mu^* \frac{s}{k^*+s}\frac{dt}{ds} ds \tag{A.7}$$

We can calculate $\frac{dt}{ds}$ from equations (A.1). First we express $x$ in $s$ and constants:

$$\begin{aligned} x(t) - x(0) &= y(s(0) - s(t)) \\ x(t) &= y(s_0 - s(t)) + x_0 \end{aligned} \tag{A.8}$$

Then we can work out $\frac{ds}{dt}$:

$$\begin{aligned} \frac{ds}{dt} &= -\frac{1}{y}\mu\frac{s}{k+s}x \\ &= -\frac{1}{y}\mu\frac{s}{k+s}(y(s_0 - s) + x_0) \\ &= -\mu\frac{s(\frac{x_0}{y} + s_0 - s)}{k+s} \end{aligned} \tag{A.9}$$

Then we get:

$$\frac{dt}{ds} = -\frac{1}{\mu} \frac{k+s}{s(\frac{x_0}{y} + s_0 - s)} \tag{A.10}$$

Now we can use equation (A.10) to calculate the integral (A.7):

$$\begin{aligned}
\int_{s_0}^{0} \mu^* \frac{s}{k^*+s} \frac{dt}{ds} ds &= \frac{\mu^*}{\mu} \int_0^{s_0} \frac{k+s}{k^*+s} \frac{1}{\frac{x_0}{y} + s_0 - s} ds \\
&= \frac{\mu^*}{\mu} \left[ \frac{k-k^*}{k^* + \frac{x_0}{y} + s_0} \log(k^*+s) - \frac{k + \frac{x_0}{y} + s_0}{k^* + \frac{x_0}{y} + s_0} \log(\frac{x_0}{y} + s_0 - s) \right]_{s=0}^{s=s_0} \\
&= \frac{\mu^*}{\mu} \left( \frac{k-k^*}{k^* + \frac{x_0}{y} + s_0} \log\left(\frac{k^*+s_0}{k^*}\right) + \frac{k + \frac{x_0}{y} + s_0}{k^* + \frac{x_0}{y} + s_0} \log\left(\frac{x_0 + y s_0}{x_0}\right) \right)
\end{aligned} \tag{A.11}$$

To calculate the growth factor of the mutant ($\frac{x^*(\infty)}{x^*(0)}$), we have to take $e$ to the power the above integral (see equation (A.6)):

$$\frac{x^*(\infty)}{x_0^*} = \left(\frac{k^*+s_0}{k^*}\right)^{\frac{\mu^*}{\mu} \frac{k-k^*}{k^* + \frac{x_0}{y} + s_0}} \left(\frac{x_0 + y s_0}{x_0}\right)^{\frac{\mu^*}{\mu} \frac{k + \frac{x_0}{y} + s_0}{k^* + \frac{x_0}{y} + s_0}} \tag{A.12}$$

Using the definition (A.3), we can describe the fitness as:

$$\frac{\log(x_\infty^*/x_0^*)}{\log(x_\infty/x_0)} = \frac{\log\left[ \left(\frac{k^*+s_0}{k^*}\right)^{\frac{\mu^*}{\mu} \frac{k-k^*}{k^* + \frac{x_0}{y} + s_0}} \left(\frac{x_0 + y s_0}{x_0}\right)^{\frac{\mu^*}{\mu} \frac{k + \frac{x_0}{y} + s_0}{k^* + \frac{x_0}{y} + s_0}} \right]}{\log\left[\frac{x_0 + y s_0}{x_0}\right]} \tag{A.13}$$

To estimate the effect of a mutant with different growth properties, we determine the expression for the invasion fitness in a fluctuating environment. The invasion fitness ($W^*$) of the mutant with a certain maximal growth rate ($\mu^*$ or $\mu^*_{max}$), substrate affinity ($k^*$ or $k^*_S$) and numerical yield ($y^*$ or $Y^*_{X/S}$) in an environment of residents (with corresponding properties $\mu$ or $\mu_{max}$, $k$ or $k_S$ and $y$ or $Y_{X/S}$) is (see Appendix section A.3 for the derivation):

$$W^* = \frac{\ln(x_\infty^*/x_0^*)}{\ln(x_\infty/x_0)} = \frac{\ln\left[ \left(\frac{k^*+s_0}{k^*}\right)^{\frac{\mu^*}{\mu} \frac{k-k^*}{k^* + \frac{x_0}{y} + s_0}} \left(\frac{x_0 + y s_0}{x_0}\right)^{\frac{\mu^*}{\mu} \frac{k + \frac{x_0}{y} + s_0}{k^* + \frac{x_0}{y} + s_0}} \right]}{\ln\left[\frac{x_0 + y s_0}{x_0}\right]} \tag{A.14}$$

When the mutant is the same as the resident ($\mu^* = \mu$ and $k^* = k$), the growth factor (relative increase in population size) of the mutant ($\frac{x^*(\infty)}{x^*(0)}$) is exactly that of the resident ($\frac{x_0 + y s_0}{x_0}$) and $W^* = 1$, as expected. When only the maximal growth rates is different ($k^* = k$ and $\mu^* \neq \mu$), the growth factor of the mutant is the growth factor of the resident to the power $\frac{\mu^*}{\mu}$ and the invasion fitness is $\frac{\mu^*}{\mu}$. The effect of a difference in substrate affinity ($k^* \neq k$) is more complex (see Section A.5), as well as the effect of a trade-off between maximal growth and affinity (see Section A.4).

Whether strains with higher affinity or with higher maximal growth rate can invade depends on the conditions, such as the initial substrate concentration ($s_0$) and the initial biomass ($x_0$). A mutant with a higher maximal growth rate ($\mu^*$) but a lower affinity (higher $k^*$) benefits from long periods of high nutrients, that is a high initial substrate ($s_0$) and a low initial biomass ($x_0$). Indeed, it follows from the above equation that a high $s_0$ leads to a relatively stronger effect of $\mu^*$ on the growth factor (and therefore a higher invasion fitness) while a high $x_0$ leads to a relatively larger effect of $k^*$ (see Section A.5).

The numerical yield is not relevant for the fitness of an invading strain, as we can scale the initial biomass ($x_0$) with the yield and eliminate the yield. The growth rate might be affected by the yield, but that effect could be added as a tradeoff between the parameters $\mu$, $k$ and $y$ (as in Figure A.5A). There is no selection on $y$, but any result that is affected by $x_0$ is inversely affected by $y$. Therefore, the yield of the resident strain does affect the fitness of other invading mutants.

##### A.3.1 Constant dilution ($d$) instead of constant initial biomass ($x_0$)

Here we do not start every transfer with a fixed biomass concentration, but instead fix the dilution from one transfer to the next. This means that the biomass at the beginning is equal to the final biomass divided by the dilution factor $d$:

$$
\begin{aligned}
x_0 &= \frac{x_\infty}{d} = \frac{x_0 + ys_0}{d} \\
x_0 &= \frac{ys_0}{d-1}
\end{aligned}
\tag{A.15}
$$

The fitness then becomes:

$$
\frac{\log(x^*_\infty/x^*_0)}{\log(x_\infty/x_0)} = \frac{\log\left[\left(\frac{k^*+s_0}{k^*}\right)^{\frac{\mu^*}{\mu}\frac{k-k^*}{k^*+\frac{d}{d-1}s_0}} d^{\frac{\mu^*}{\mu}\frac{k+\frac{d}{d-1}s_0}{k^*+\frac{d}{d-1}s_0}}\right]}{\log d}
\tag{A.16}
$$

In this case the yield is removed from the equation, because $x_0$ will automatically be adjusted by the yield, and the nutrient profile will not depend on the yield anymore.

#### A.4 Trade-offs between $\mu_{max}$, $k_S$ and $y$

##### A.4.1 *E. coli* from experimental data

A trade-off between maximal growth rate and affinity has been fitted to experimental data[6]:

$$
\mu_{max} = \mu^{ref}\frac{log(\frac{k}{k^{ref}})}{log(\frac{k}{k^{ref}})+1}
\tag{A.17}
$$

There was a small issue with the parameters reported in the paper (personal communication) and we have converted the parameter $k^{ref}$ from $\mu g\,L^{-1}$ to $mM$. This results in the parameters $\mu^{ref} = 1.23\,h^{-1}$ and $k^{ref} = 3.056 \cdot 10^{-5}\,mM$. Since we have no information on the yield for this data, we used a constant yield derived from the LTEE of the Lenski lab of $474 \cdot 10^9$ cells $\text{mmol}^{-1}$[5].

##### A.4.2 *E. coli* from kinetic model of central metabolism

For an actual evolutionary trade-off, we need to find the pareto optimum between different properties. Maximal growth rates and affinities in experimental conditions are usually for cells that are optimized for one of these properties. Another way to approach optimal conditions is by making a computational model of a species and then optimizing the model for different growth properties. We used a model of the central carbon metabolism of *E. coli*[7] for this purpose. Optimal yield and growth under defined external conditions are achieved by elementary flux modes[8], allowing us to find the pareto front by selecting the pareto optimal EFMs. Fitting a function through these pareto optimal EFMs for $\mu_{max}$ and $k_S$, and another function for $Y_{X/S}$ and $\mu_{max}$ leads to:

$$
\begin{aligned}
\mu_{max} &= \mu^{ref}\frac{(k^{ref})^n}{\frac{1}{k}^n + (k^{ref})^n} \\
Y_{X/S} &= y_{max}\frac{\mu_{max}}{\mu_{max} + y_{ref}}
\end{aligned}
\tag{A.18}
$$

With $\mu^{ref} = 0.8$, $k^{ref} = 417$ and $n = 2.7$ for the first equation and $y_{max} = 495$ and $y_{ref} = 0.0726$ for the second equation. For anaerobic growth we observe a different shape trade-off:

$$
\mu_{max} = \mu_1^{ref}\frac{k}{k + k_1^{ref}} + \mu_2^{ref}\frac{k^n}{k^n + (k_2^{ref})^n}
\tag{A.19}
$$

With $\mu_1^{ref} = 0.1$, $k_1^{ref} = 3.9 \cdot 10^{-3}$, $\mu_2^{ref} = 0.21$, $k_2^{ref} = 2.3 \cdot 10^{-2}$ and $n = 5.6$. Under anaerobic conditions we observed a relatively constant yield of $40 \cdot 10^9$ cells $\text{mmol}^{-1}$. See Fig. A.1 for the fits.

##### A.4.3 *S. cerevisiae* from experimental data

We used data from *S. cerevisiae* with all glucose transporters knocked out and single transporters or chimeras of two different transporters placed back[9]. The $K_M$'s of the transporters as well as the growth rates are reported for these strains. We checked with a model of *S. cerevisiae* glycolysis[10] and when the transporter $K_M$ is the only difference, this also reflects the substrate concentration of half maximal glycolysis rate. Assuming glycolysis rate is related to growth rate, in this case the $K_M$ of the transporter will be a good proxy for the $K_S$ of the cell. The function for these data is:

$$
\begin{aligned}
\mu_{max} &= \mu^{ref}\frac{k_S}{k_S + k^{ref}} \\
Y_{X/S} &= y_{max}\frac{y_{ref}^m}{\mu_{max}^m + y_{ref}^m}
\end{aligned}
\tag{A.20}
$$

With $\mu^{ref} = 0.34$, $k_S = 0.77$, $y_{max} = 7$, $y_{ref} = 0.3$ and $m = 10$. The fits are depicted in Figure A.2.

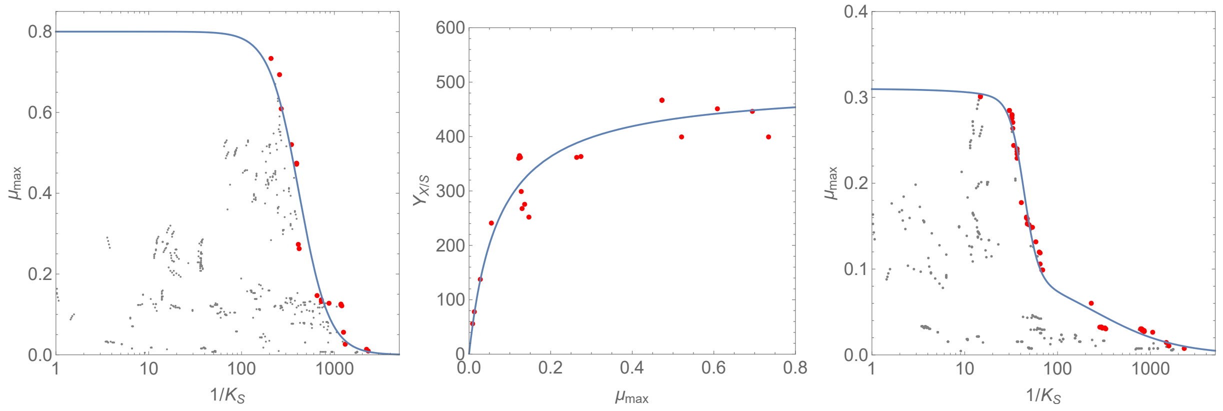

Figure A.1: **Trade-off curves fitted to the pareto front for a computational model of *E. coli*.** A computational model is used to calculate $\mu_{max}$ and $k_S$ of all Elementary Flux Modes (dots). The modes that form the pareto front (no other modes exist with both higher $\mu_{max}$ and $k_S$) are colored red, and a function is fitted through those values (Eqs (A.18) and (A.19)). **Left:** The fit of the trade-off curve between $\mu_{max}$ and $k_S$ under aerobic conditions. **Middle:** The fit of the trade-off curve between $\mu_{max}$ and $Y_{X/S}$ under aerobic conditions. **Right:** The fit of the trade-off curve between $\mu_{max}$ and $k_S$ under anaerobic conditions.

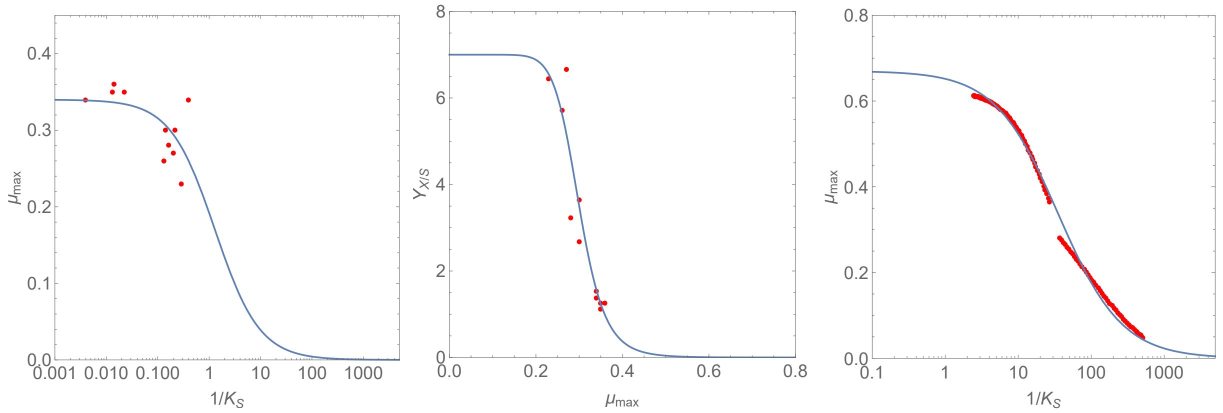

Figure A.2: **Trade-off curves fitted to experimental data and a computational model of *S. cerevisiae*.** Data from chimeric transporters is used to generate strains with different properties[9] (dots). **Left:** A curve is fit for the $\mu_{max}$-$k_S$ trade-off (eq (A.20)). **Middle:** A curve is fit for the $\mu_{max}$-$Y_{X/S}$ tradeoff (eq (A.20)). **Right:** A curve is fit to a self-replicator model parameterized on *S. cerevisiae*[11] (eq (A.21)).

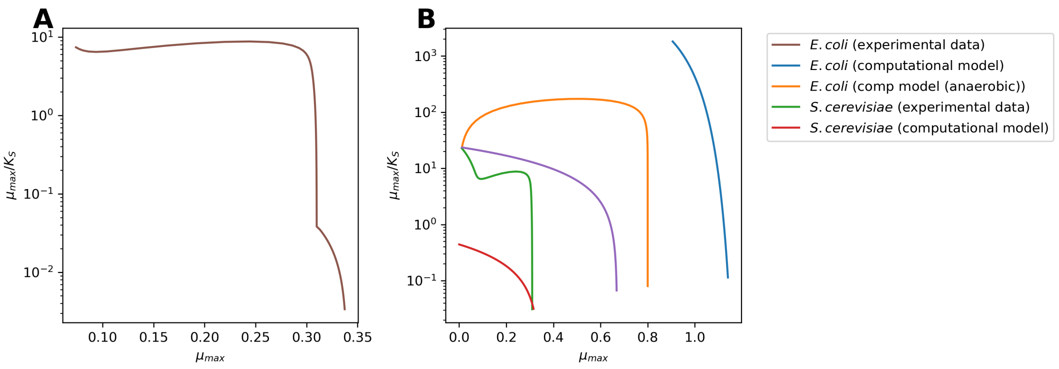

Figure A.3: Trade-offs between growth parameters. **A** Trade-off between $\mu_{max}$ and $\mu_{max}/k_S$ for the two-species case. **B** Trade-off between $\mu_{max}$ and $\mu_{max}/k_S$ for the individual curves.

##### A.4.4 *S. cerevisiae* from self-replicator model

We also investigated a computational approach to get a trade-off curve for *S. cerevisiae*. We used a self-replicator model that includes all reactions to replicate itself, although in a very simplified form, and is parameterized on experimental data[11]. When we optimize this self-replicator model on different levels of the resource (glucose), and then fit Monod curves through each of the optimized parameters, we obtain different values for $\mu_{max}$ and $k_S$. We fitted a trade-off curve through the corresponding $\mu_{max}$ and $k_S$ values:

$$\mu_{max} = \mu^{ref} \frac{k_S}{k_S + k^{ref}} \tag{A.21}$$

with $\mu^{ref} = 0.67$ and $k^{ref} = 0.028$. See Figure A.2 for the fit. In the computational results, *S. cerevisiae* either fully respires or fully ferments. The yield is constant for such a regime, and therefore the yield that is used is a step function:

$$\begin{aligned} Y_{X/S} &= 7.8 \ \text{ when } \mu_{max} < 0.3 \\ Y_{X/S} &= 3.7 \ \text{ when } \mu_{max} \geq 0.3 \end{aligned} \tag{A.22}$$

To convert the yield from gramDW mmol Glucose$^{-1}$ to $10^9$ cells mmol Glucose$^{-1}$ we used a value of $15 \cdot 10^{-12}$ gram per cell.

##### A.4.5 Trade-off curves in $\mu_{max}$-$\mu_{max}/k_S$ space

For the $\mu_{max}$-$k_S$ trade-off to be effective, the species with a higher $\mu_{max}$ needs to have a lower $\mu_{max}/k_S$, because this is a minimum requirement for having a higher growth rate at low nutrient concentrations, as also mentioned in[12]. Those curves are depicted in figure A.3.

##### A.4.6 Conversion to a normalized substrate specialization parameter

When integrating the trade-off in our models we have chosen a 'specialization parameter' $u$, that runs from 0 (specialization on low resource) to 1 (specialization on high resource). The general conversion is (Figure figure A.5):

$$k = k_{min} 10^{u \log_{10}\left(\frac{k_{max}}{k_{min}}\right)} \tag{A.23}$$

##### A.4.7 Combined trade-off of two species

To investigate evolutionary coexistence of two species while still applying the same analytical and graphical analyses as for the single species case, we can make a single trade-off curve that encompasses the two separate trade-off curves. Because mutations cannot change one species in another, we have to make sure there are no mutations across the species boundary, which is likely if we have the species boundary in a fitness minimum.

To apply this we have combined the experimentally derived trade-off for *S. cerevisiae* (section A.4.3) and the computationally derived trade-off for *E. coli* for anaerobic conditions (section A.4.2). We used equation (A.19) below $k_S = 8.21$ and equation (A.20) above (see Figure 4).

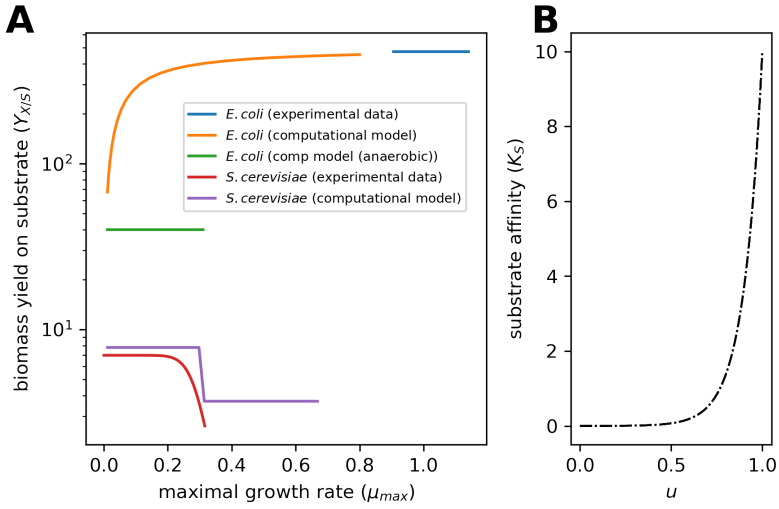

Figure A.4: **A** All relations between $\mu_{max}$ and yield. There is no direct selection on the yield, but the yield affects the dynamics of substrate depletion, and therefore the environment for other species. There is quite some literature on trade-offs between $\mu_{max}$ and $Y_{X/S}$ (see Beardmore et al. [13] for a summary), but here we used yields from the same sources as the other growth parameters as much as possible. **B** Conversion to normalized substrate specialization parameter $u$ for $k_{min} = 0.0005$ and $k_{max} = 10$ (Equation (A.23)).

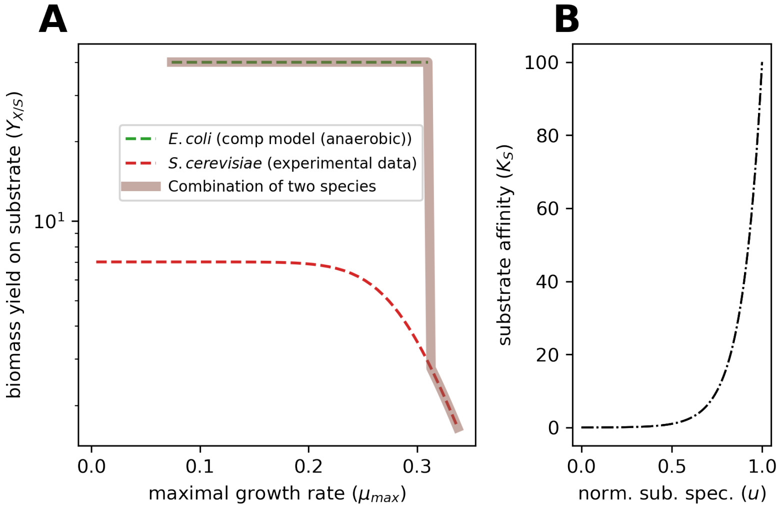

Figure A.5: **A** Link between $\mu_{max}$ and yield for the two-species case. **B** Conversion to normalized substrate specialization parameter $u$ for $k_{min} = 0.01$ and $k_{max} = 100$ that was used for the two-species simulations (Equation (A.23)).

#### A.5 Selection gradient

The selection gradient $S_X$ for a property $X$ is the proportional partial derivative of the invasion fitness to the property. The selection coefficient (eq (A.4)) for $\mu$ t is always equal to 1, because eq (A.14) is proportional to $\mu^*$, the derivative $\partial W/\partial\mu^*$ is exactly $W/\mu^*$. Since we assume the mutant has not influence on the nutrient profile, the selection gradient for $y^*$ is 0. For $k^*$ this selection gradient in the point $k = k^*$ is equal to (we are keeping everything but the $k$ constant, so $\mu^*/\mu = 1$). This leads to the following selection gradients:

$$\begin{aligned} S_\mu &= \frac{\mu^*}{W^*}\frac{\partial W^*}{\partial \mu^*}(\mu^* = \mu) = 1 \\ S_y &= \frac{y^*}{W^*}\frac{\partial W^*}{\partial y^*}(y^* = y) = 0 \\ S_k &= \frac{k^*}{W^*}\frac{\partial W^*}{\partial k^*}(k^* = k) = \frac{-k\left(1 + \frac{\log\left(\frac{k+s_0}{k}\right)}{\log\left(\frac{x_0+ys_0}{x_0}\right)}\right)}{k + \frac{x_0}{y} + s_0} \end{aligned} \tag{A.24}$$

Using this equation, we can calculate selection on the Monod constant in an experimental setting. For the Long Term Evolution Experiment (values from[5]) we find $S_k = -0.00657233$, which agrees with the $-0.0066$ mentioned in the paper, where they obtain this result with numerical calculations. As this selection gradient is very low (compared to $S_\mu = 1$)—and indeed they do not observe any increase in the affinity over the course of evolution —, we wondered if we expect to see selection on the Monod constant for different strains for *E. coli* or under different conditions. Higher values for $k$ lead to a higher selection gradient, and the $k$ for some *E. coli* strains is a factor 100 higher than the value of 0.727 $\mu g/mL$ reported for REL606[5]. At those higher values we can see that the selection of the $k_S$ does become important, as well as under different experimental conditions (see Figure A.6). Selection on the $k_S$ might be more important for other substrates or other organisms, e.g. if uptake is not active but based on a diffusion gradient such as for the yeast *Saccharomyces cerevisiae* the $k_S$ is usually (much) higher.

The landscape of invasion fitness and the gradient of this landscape (the selection gradient) can be visualized (Fig. A.7). $S_k$ is always lower or equal to $S_\mu$, which has implications for a $k/\mu$ trade-off, namely that the trade-off is only relevant if a 1% effect on the maximal growth rate has a bigger than 1% effect on the $k_s$.

When we plot the selection coefficient and the trade-off function for a resident in the LTEE of the Lenski lab[5] we expect the selection to be almost completely on increasing the maximum growth rate. However, when we would repeat the experiment with a 100 times lower starting glucose concentration, we would expect a much more pronounced selection on the affinity (Figure A.7).

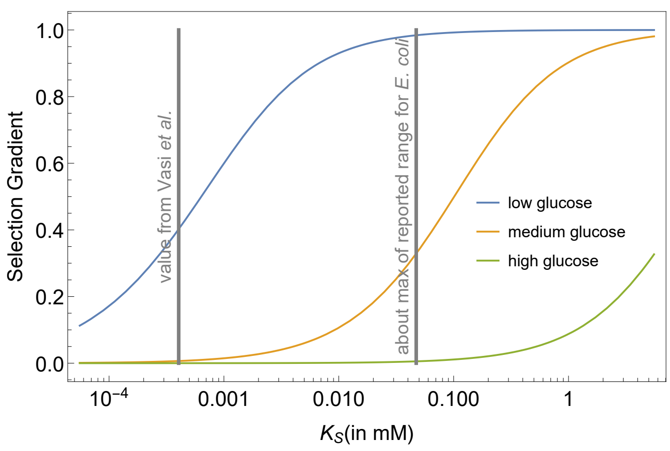

Figure A.6: Dependency of the selection gradient on the $k_s$. We have taken the selection gradient for a decrease in $k$ (minus the selection gradient). Range of $k_S$ values taken from[14]. Medium glucose is 25 $\mu$gmL$^{-1}$ = 0.1375 mM (values used in[5]), low glucose is 100 times lower, and high glucose 100 times higher.

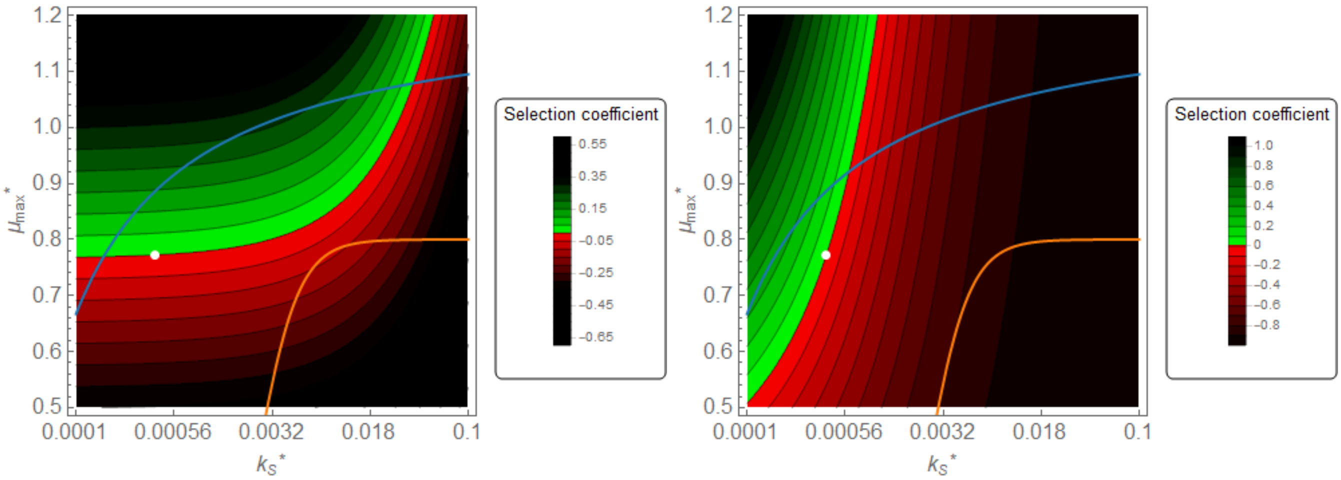

Figure A.7: **Landscapes of selection coefficients for a given resident.** The selection gradient is the slope in this landscape. Blue line is the trade-off from experimental data [6] and orange line the trade-off from computational data [7]. The resident is chosen from measurements [5] (these are better than prediction from the computational trade-off). **Left** Conditions from the LTEE of the Lenski lab [5]. Selection mostly promotes maximum growth rate (higher selection coefficients for higher $\mu_{max}$ within the trade-off curves, so mostly mutants with higher $\mu_{max}$ can invade). **Right** 100 times lower glucose conditions. Now, selection is also on affinity, even strategies with lower $\mu_{max}$ and higher $k_S$ can invade.

#### A.6 Invasive potential for different species, trade-offs and experimental conditions

When we focus only on the trade-off line, we can simplify the figures from Figure A.7 to one dimension. We can use the second dimension to plot this not only for a single resident, but for all possible residents. For the different tradeoffs (Section A.4), we can plot the pairwise invasibility plots (Figure A.8) and by mirroring them over the diagonal and overlaying them, we can show the area of ecological coexistence (Figure A.9).

In the mutual invisibility plot, we can study the evolutionary dynamics. A common way is to use a fitness function with two residents, but in this case it is not possible to derive such a function. Therefore we use a graphical approach expained in the appendix of Geritz et al. [15]. The method uses information from the shape of the pairwise invasibility plot and the mirrored pairwise invasibility plot to find where the isoclines cross the border of the area of coexistence (see Figure A.10). With that information we can conclude that there is no coexistence in the mutual invasibility plots of the layout of Figure A.9, but there is in the case of the two species together (main text Figure 4C).

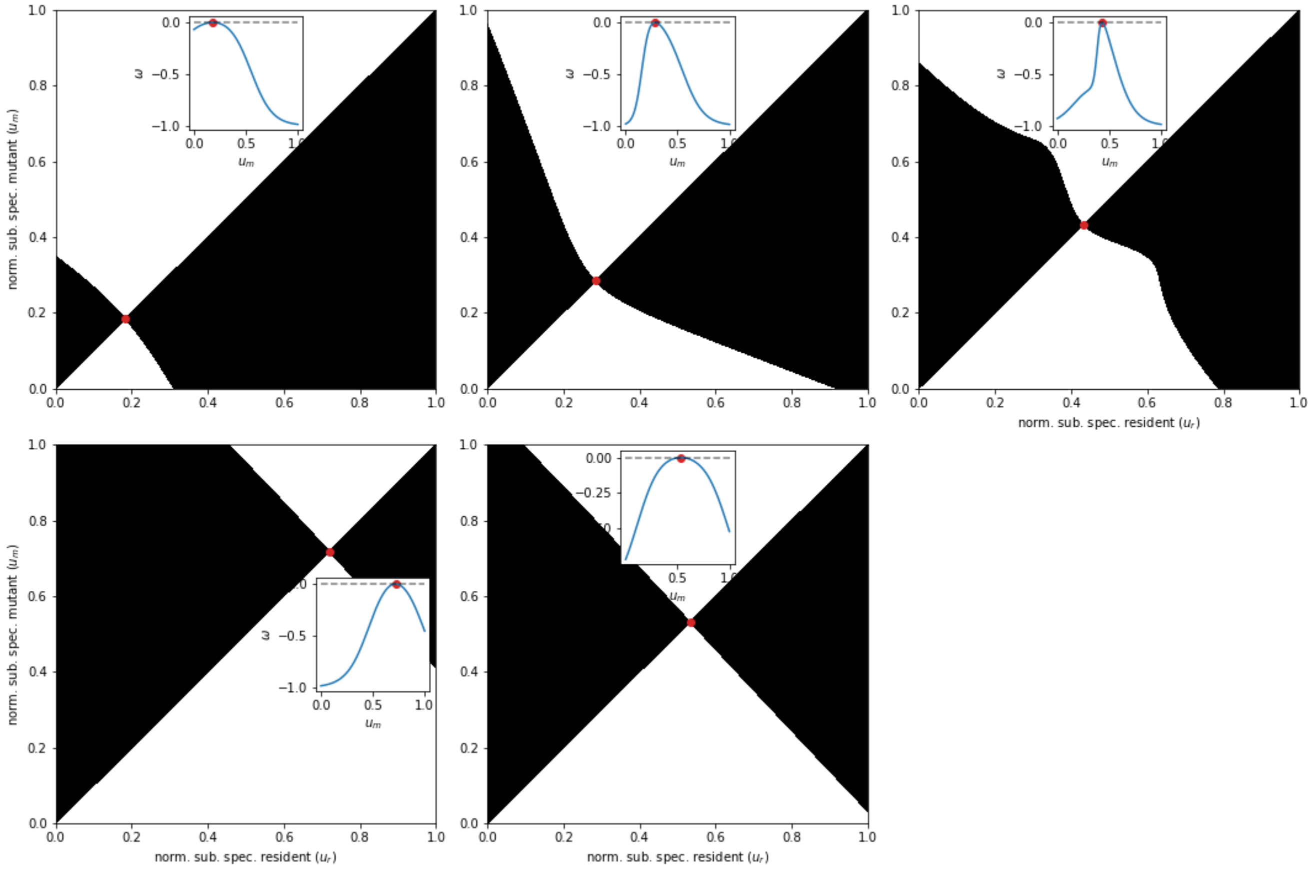

Figure A.8: **Pairwise invasibility plots for the different trade-offs Top row:** different trade-offs for *E. coli* (experimental fit, computational fit and computational fit anaerobic) with experimental conditions: initial glucose = 0.1375 mM and initial cells are $0.5{\cdot}10^6$ cells/ml. For *E. coli* $k_S$ runs from 0.1 to 10 for $u$ from 0 to 1 (see Figure A.5B and Eq (A.23)). **Bottom row:** different trade-offs for *S. cerevisiae* (experimental fit and computational fit) with experimental conditions: initial glucose = 100 mM and initial cells are $0.1{\cdot}10^6$ cells/ml. For *S. cerevisiae* $k_S$ runs from 0.1 to 100 for $u$ from 0 to 1 (see Figure 4B and Eq (A.23)).

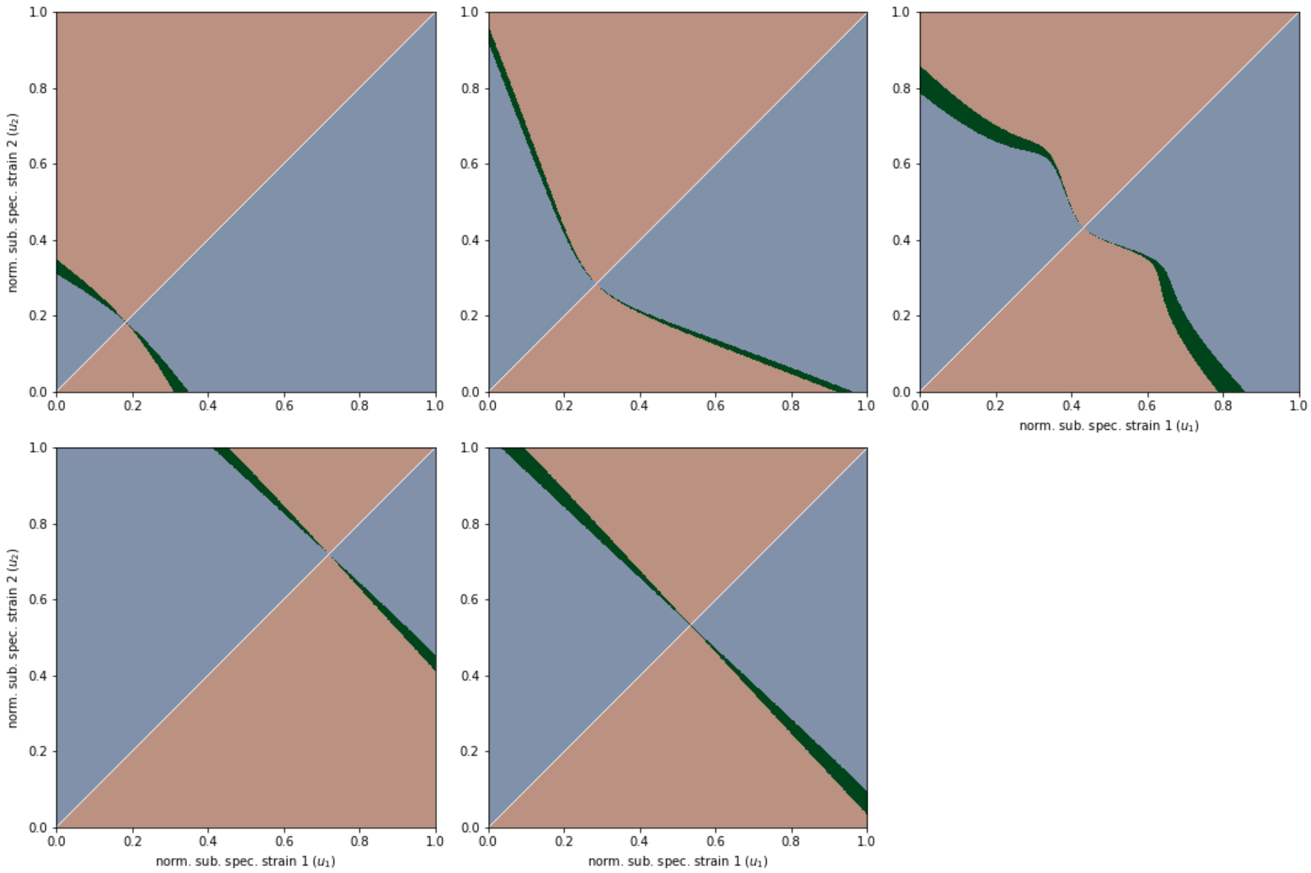

Figure A.9: **Areas of coexistence for the different trade-offs Top row:** different trade-offs for *E. coli* (experimental fit, computational fit and computational fit anaerobic) with experimental conditions: initial glucose = 0.1375 mM and initial cells are $0.5{\cdot}10^6$ cells/ml. For *E. coli* $k_S$ runs from 0.1 to 10 for $u$ from 0 to 1 (see Figure A.5B and Eq (A.23)). **Bottom row:** different trade-offs for *S. cerevisiae* (experimental fit and computational fit) with experimental conditions: initial glucose = 100 mM and initial cells are $0.1{\cdot}10^6$ cells/ml. For *S. cerevisiae* $k_S$ runs from 0.1 to 100 for $u$ from 0 to 1 (see Figure 4B and Eq (A.23)).

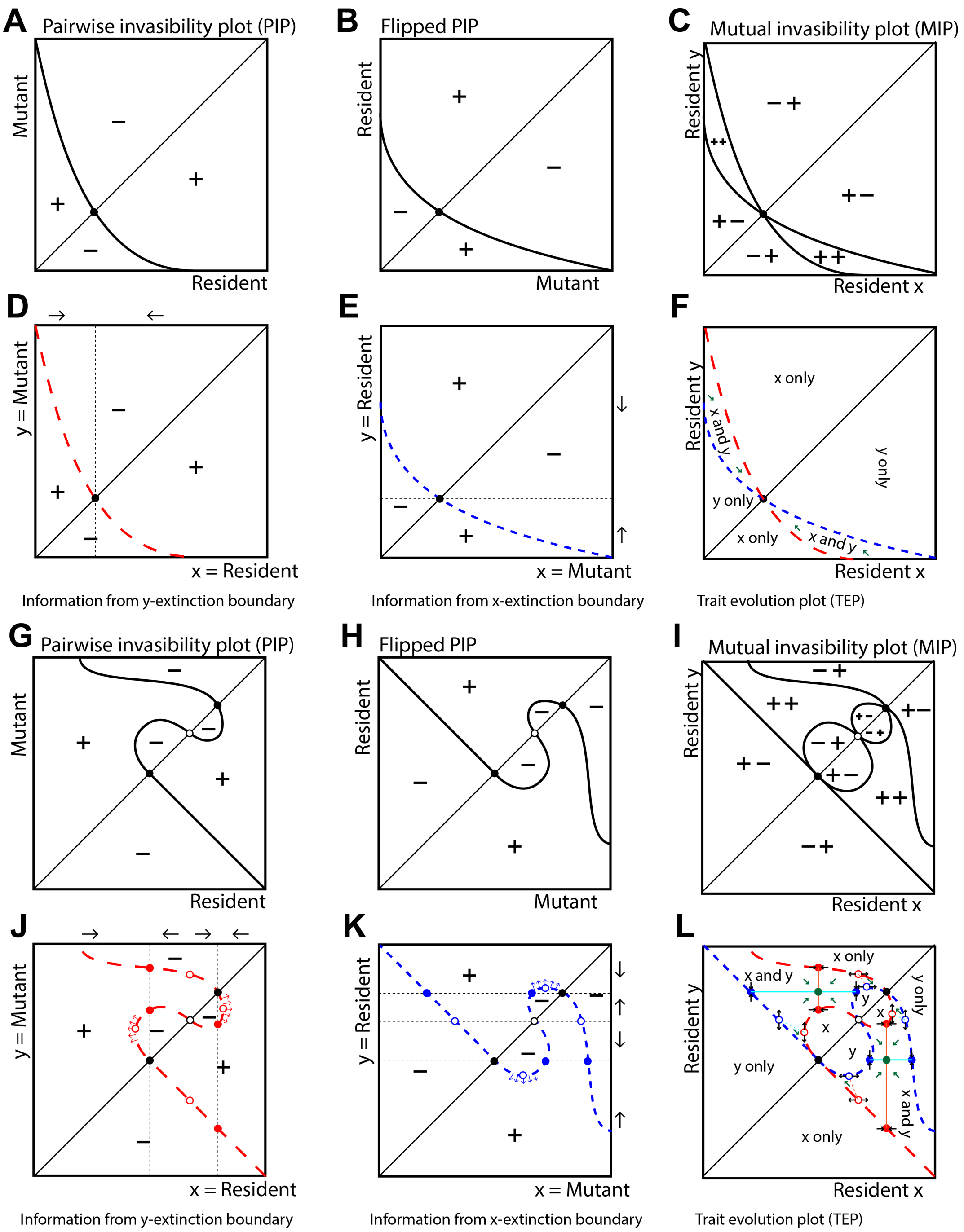

Figure A.10: **Schematic analysis of evolutionary dynamics with two residents** The two residents are labeled $x$ and $y$. **A-F** Analysis of the dynamics belonging to a species with a 'simple' trade-off (main text Figure 2A). Overlaying the PIP (**A**) and the PIP flipped over the diagonal (**B**) leads to the area of ecological coexistence (**C**). On the line originating from the PIP, $y$ will co extinct, because this is the border between the ++ and the +- area (**D**). The arrows above shows the behaviour with only $x$ present (which can be read from the diagonal); an arrow to the right means higher types can invade and to the left that lower types can invade. Close to the extinction boundary these evolutionary dynamics will be the same in the $x$-direction in the coexistence boundary. Analysis on the flipped PIP gives the same analysis on the $x$ extinction boundary (**E**). Putting this information together we can visualize the dynamics in the area of coexistence (green arrows in **F**). **G-L** Analysis for the dynamics belonging to a species with a 'bi-phasal' trade-off (main text Figure 2C). The analysis is similar, but now there are intersections from the vertical lines through the singular points in the PIP with the $y$-extinction boundary, which signify where the $x$-isoclines cross the $y$ extinction boundary (**J**). For stable singular points this leads to an intersection of a stable isocline (closed circles) and unstable singular points create an intersection of an unstable isocline (open circles). Moreover, the selection gradient for $y$ has to be perpendicular to the $y$ extinction boundary, and therefore there is a switch in direction for the extreme points in the $x$-direction. Because of the shape of the extinction boundary, these are intersection points of unstable isoclines. Similar analysis on the flipped PIP (**K**) gives more intersection points, which are connected in the trait evolution plot (**L**). Although the shape of the isoclines in the interior of the coexistence area is unclear, it is sure that stable $x$ and $y$ iscolines must intersect (orange and blue solid lines), which leads to a stable state with two residents, called an evolutionary stable coalition (green dot).

#### A.7 Stochastic simulations

In the case of multiple resident species or strains (we will call them types), or when we explicitly model several mutants with stochastic simulations, the model system for $n$ types becomes:

$$\begin{aligned}\frac{dx_i}{dt} &= \mu_i \frac{s}{k_i + s} x_i \\ \frac{ds}{dt} &= -\sum_{i=1}^{n} \frac{1}{y_i} \mu_i \frac{s}{k_i + s} x_i\end{aligned} \tag{A.25}$$

We have used the two-species trade-off to determine $\mu_{max}$ and $Y_{X/S}$ from $u$ (Section A.4.7 and Figure 4). For the simulations, we allowed for 51 different phenotypes at the same time, and tracked their fractions at the beginning and end of a fluctuation to determine the relative growth rate of a phenotype. All growing phenotypes could generate offspring with small but stochastic mutations, with a small chance, whenever free phenotypes were available (code adapted from online available code by Guilhem Doulcier). We simulated these systems until a stable phenotype distribution was reached (Figure A.11).

To plot the phenotype trajectories in Figure 4C, we needed to convert the collection of phenotypes to one or two main trait values. We did this by taking only the phenotypes which had a density of more than 1/1000th of the most abundant phenotype. Then we checked if all those phenotypes were close together or far apart, and created either one or two groups. For every group we took the weighted mean to determine the consensus phenotype. See Figure A.12 for two examples of the outcome of this algorithm.

The evolutionary stable consortium was taken as the endpoint of the simulations starting from two phenotypes (A.11C), and then calculating the consensus phenotypes for that endpoint. To check if this consortium is indeed evolutionarily stable, we numerically calculated the invasion fitness of all possible strategies in the consortium (see inset in Figure 4C).

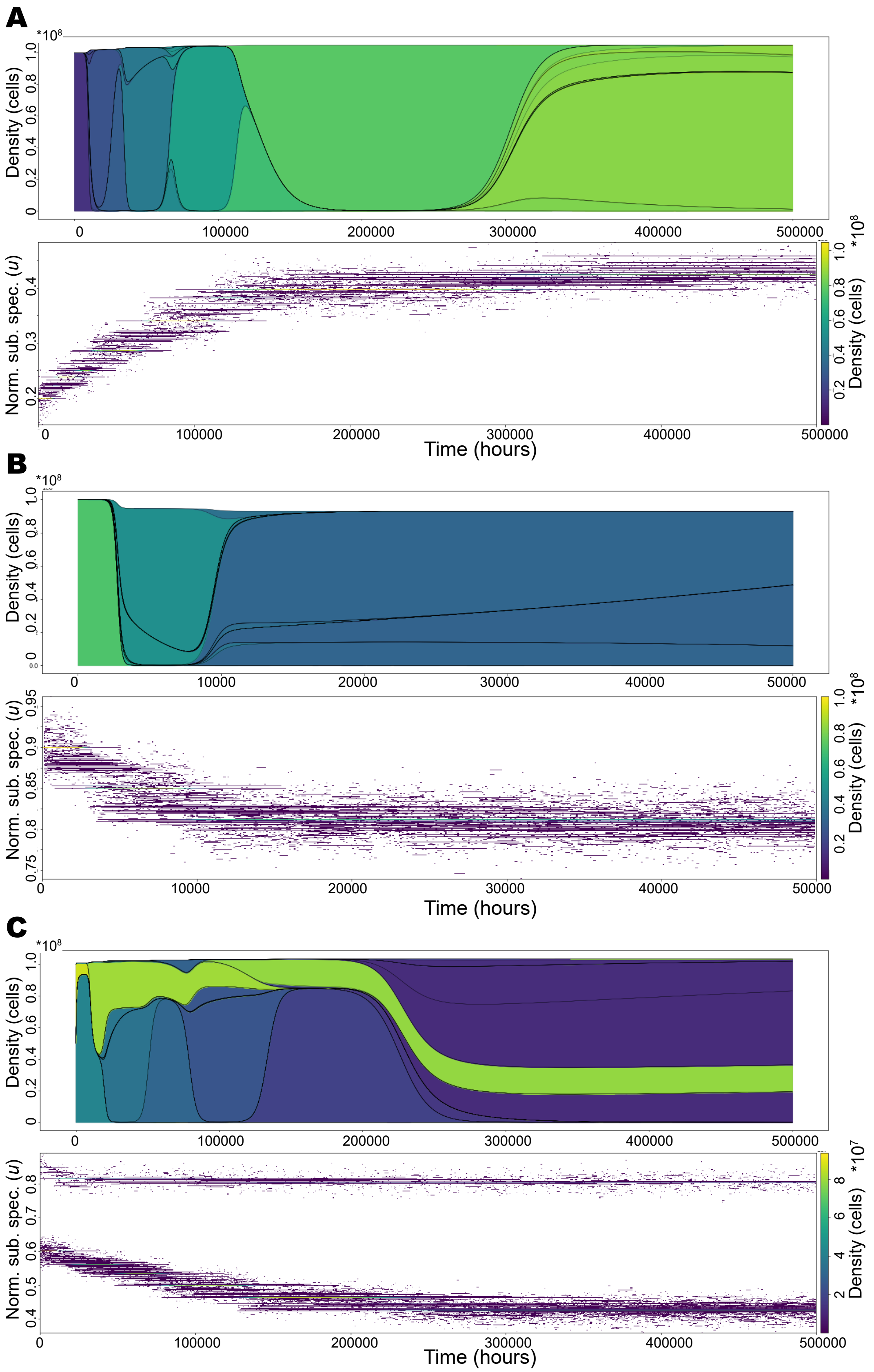

Figure A.11: **Trajectories of stochastic simulations starting from different initial conditions. A** and **B** Simulations start from a single initial phenotype, a low (**A**) and high (**B**) substrate specialist, to end up at a single species ESS (see Figure 4C). **C** Simulation starts from a polymorphic population with two phenotypes, and ends up in the evolutionary stable consortium (see Figure 4C).

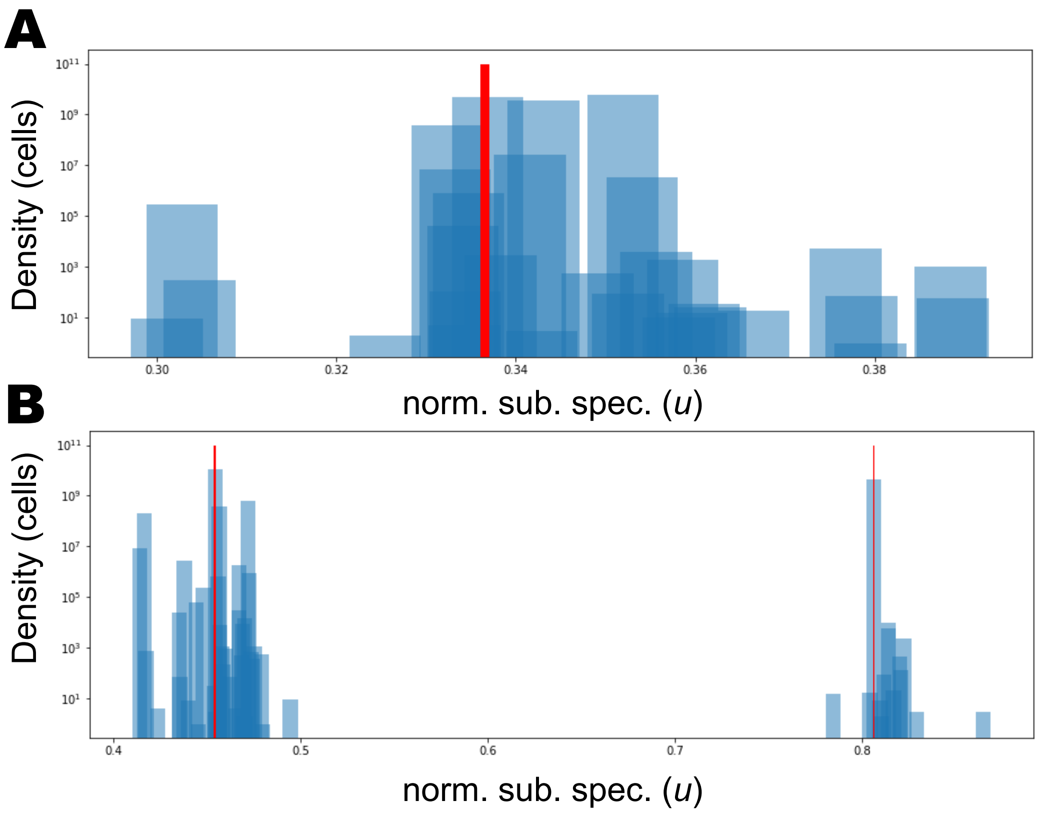

Figure A.12: The red line(s) show(s) the traits used for plotting the trajectories in Figure 4C for two different examples: **A** with one dominant type and **B** with two types.
